# The lipid raft-linker gene *Raftlin-2* is expressed in migrating neural crest cells

**DOI:** 10.64898/2026.06.12.731942

**Authors:** Mallorie P. Jenne, Ilya Grabylnikov, Michael L. Piacentino

## Abstract

Transient plasma membrane domains called lipid rafts have emerged as important regulators of signal transduction. These territories are formed by lipid-lipid and lipid-protein interactions, and these local interactions can be scaffolded by resident lipid raft organizing protein family members. While roles for lipid rafts have been described for multiple signaling pathways in many contexts, their *in vivo* prevalence and role during embryonic development remains incompletely understood. Here we examined gene expression for the Raftlin family of lipid raft organizing proteins, Raftlin (*RFTN1*) and Raftlin-2 (*RFTN2*), over the course of early vertebrate development, with a focus on neural crest cell dynamics. By analyzing transcriptomic data across vertebrate species, we identified conserved patterns of *RFTN1* and *RFTN2* expression across species, where *RFTN1* is broadly expressed at low levels, while *RFTN2* is distinctly enriched in neural crest cells. We used fluorescent *in situ* hybridization to spatially define Raftlin gene expression patterns in the early avian embryo. Our results show that *RFTN1* is broadly expressed with periods of enrichment in the developing paraxial mesoderm. In contrast, *RFTN2* expression is strongly enriched in neural crest cells, beginning during specification and persisting through migration, with additional expression in both the cranial and intermediate mesoderm. Together, these patterns suggest that Raftlins may play important roles in regulating signaling during development with specific roles in somitogenesis and in neural crest and mesodermal cell migrations.

## Introduction

Lipid rafts are nanoscale, transient membrane domains enriched in sphingolipids and cholesterol that often facilitate the enrichment or exclusion of critical signaling components, thereby acting as adaptable signaling platforms (Isik and Cizmecioglu, 2023; Roy and Patra, 2023; Sezgin et al., 2017; Simons and Ikonen, 1997). Accordingly, lipid rafts within the plasma membrane have been implicated in regulating numerous signaling pathways that play key roles in development (Roy and Patra, 2023); however, the scope and prevalence of lipid raft function during embryogenesis remains unclear. While lipid rafts form due to transient lipid-lipid and lipid-protein interactions, their stability is regulated by families of lipid raft scaffolding proteins. Thus, insight into their involvement in developmental processes may come from carefully interrogating the spatiotemporal genes expression profiles of such lipid raft scaffolding proteins.

To this end, here we report the expression of two such lipid raft organizers over the course of vertebrate embryogenesis with a particular focus on neural crest cell development. Neural crest cells are a multipotent population of embryonic cells in the vertebrate embryo that contribute to many different adult tissues including the craniofacial skeleton, neurons and glia of the peripheral nervous system, septation in the heart, and melanocytes in the skin (Stundl et al., 2026; Vega-Lopez et al., 2018). During their development neural crest cells make complex specification and cell fate decisions, undergo coordinated proliferation, epithelial-to-mesenchymal transition (EMT), and extensive migration (Piacentino et al., 2020; Shellard and Mayor, 2019; Stundl et al., 2026). These behaviors require carefully regulated levels of developmental signals including those of the Wnt, Fibroblast Growth Factor (FGF), Bone morphogenetic protein (BMP), Transforming growth factor β (TGF-β), and Notch pathways (Prasad et al., 2019; Stundl et al., 2026). Since lipid rafts have been implicated in regulating each of these signaling pathways (Roy and Patra, 2023), neural crest cells may serve as a tractable *in vivo* model for studying the role of lipid rafts in a wide range of cellular processes.

Previous studies have shown expression of several lipid raft organizing proteins to be enriched in neural crest cells, suggesting that lipid rafts play essential roles at specific times or in specific subpopulations of developing neural crest. Notably, a group of raft organizing proteins called Flotillins (also known as Reggies) are expressed in neural crest cells and neural crest cell derivatives in zebrafish and *Xenopus* embryos (Langhorst et al., 2005; Pandur et al., 2004; von Philipsborn et al., 2005). More recent work analyzing differential gene expression across neural crest EMT has shown upregulated expression of the another lipid raft organizing protein, *RFTN2* (Raftlin-2), during avian cranial neural crest migration (Piacentino et al., 2022). These expression profiles raise the possibility that lipid raft function, regulated by these organizing proteins, serves an important role in regulating essential cellular processes during neural crest development.

Raftlin proteins, named for lipid **<u>raft</u>**-**<u>lin</u>**king functions, are thought to act as scaffolds that localize to and regulate cell signaling on the plasma membrane (Lin et al., 2025; Matsumoto and Tatematsu, 2016). Raftlin (gene symbol *RFTN1*), and its paralog Raftlin-2 (gene symbol *RFTN2*), were first identified as necessary for T- and B-cell receptor signaling by concentrating receptor and downstream transmission networks in lipid raft domains (Matsumoto and Tatematsu, 2016; Saeki et al., 2009, 2003). Raftlin proteins localize to the plasma membrane by reversible acylation on N-terminal glycine and cysteine residues, and this membrane targeting strategy is required to mediate the interactions between B- and T-cell signaling components (Saeki et al., 2009, 2003). Despite lack of overt developmental phenotypes in Raftlin and Raftlin-2 single and double knockout mice (Saeki et al., 2009; Tatematsu et al., 2016), their protein products have since been shown to function beyond T- and B-cell receptor signaling with essential functions in cell proliferation, survival, and even migration in other contexts. For example, Raftlin promotes proliferation through the AKT/p38 Mitogen-Activated Protein Kinase (MAPK) signaling pathway in gastric cancer (Deng et al., 2022), and *RFTN2* expression correlates with increased metastasis in B16 melanoma lines (Kienzler et al., 2025). On the other hand, Raftlin negatively regulates endothelial cell migration by stabilizing vascular endothelial growth factor receptor 2 (VEGFR2) with its co-receptor neuropilin-1 on the plasma membrane and hinders their endocytosis, thereby modulating the downstream effects of VEGF signaling and diminishing focal adhesion engagement (Bayliss et al., 2020). This finding is particularly relevant to neural crest biology, since neural crest migration requires chemoattraction toward chemokines such as VEGF (McLennan et al., 2010; Wiszniak et al., 2015). Together these observations suggest that Raftlin proteins may play important roles in neural crest cell development by tuning essential signaling events.

To better understand the possible roles of lipid raft-organizing proteins during neural crest cell development, here we carefully characterized the dynamic spatiotemporal expression pattern of the Raftlin genes *RFTN1* and *RFTN2* during early vertebrate development. Drawing upon previous transcriptomics studies, we show evidence that *RFTN2* expression, but not *RFTN1*, is conserved in neural crest cells across amniotes. We then use fluorescent *in situ* hybridization in early avian embryos to define their spatiotemporal expression pattern at multiple stages of vertebrate development that encompass neural crest cell formation, migration, and fate decision. We show that while *RFTN2* is highly expressed in specified and migratory neural crest cells along multiple axial levels, strong *RFTN1* expression is instead restricted to the developing paraxial mesoderm. These findings implicate *RFTN2* as a potential regulator of neural crest cell development, while *RFTN1* instead may regulate cell signaling in other lineages.

## Results and Discussion

### Raftlin gene expression is dynamic during neural crest development across multiple species

To determine if Raftlin proteins may play a role in important developmental transitions, we sought to characterize their expression patterns in neural crest cells, a highly tractable *in vivo* model of several key developmental processes. To this end, we examined Raftlin gene expression in published transcriptome datasets from multiple amniote species including chick and mouse embryos, and embryonic stem cell (ESC)-derived human neural crest-like cells. First, we examined bulk RNA sequencing results collected from whole chicken embryos, as well as cranial neural crest cells sorted by *TFAP2AE1::GFP* expression, across multiple stages of development from Hamburger Hamilton stages HH6 through HH16 (Hovland et al., 2022). We plotted normalized expression levels for *RFTN1* and *RFTN2*, together with *SOX10* to display phases of neural crest specification and migration (Fig. 1A). The results show relatively stable expression of *RFTN1* across development, both in the whole embryo and in the neural crest samples. Conversely, *RFTN2* expression becomes prominently upregulated in neural crest cell samples following the onset of *SOX10* expression, which is maintained from HH8 through HH16, with a slight downregulation by HH14 (Fig. 1A). This phase of peak expression coincides with the timeframe during which cranial neural crest cells are migrating most extensively through the head, suggesting that *RFTN2* expression is regulated alongside the neural crest specification and migration gene regulatory networks (Stundl et al., 2026).

**Fig 1.**
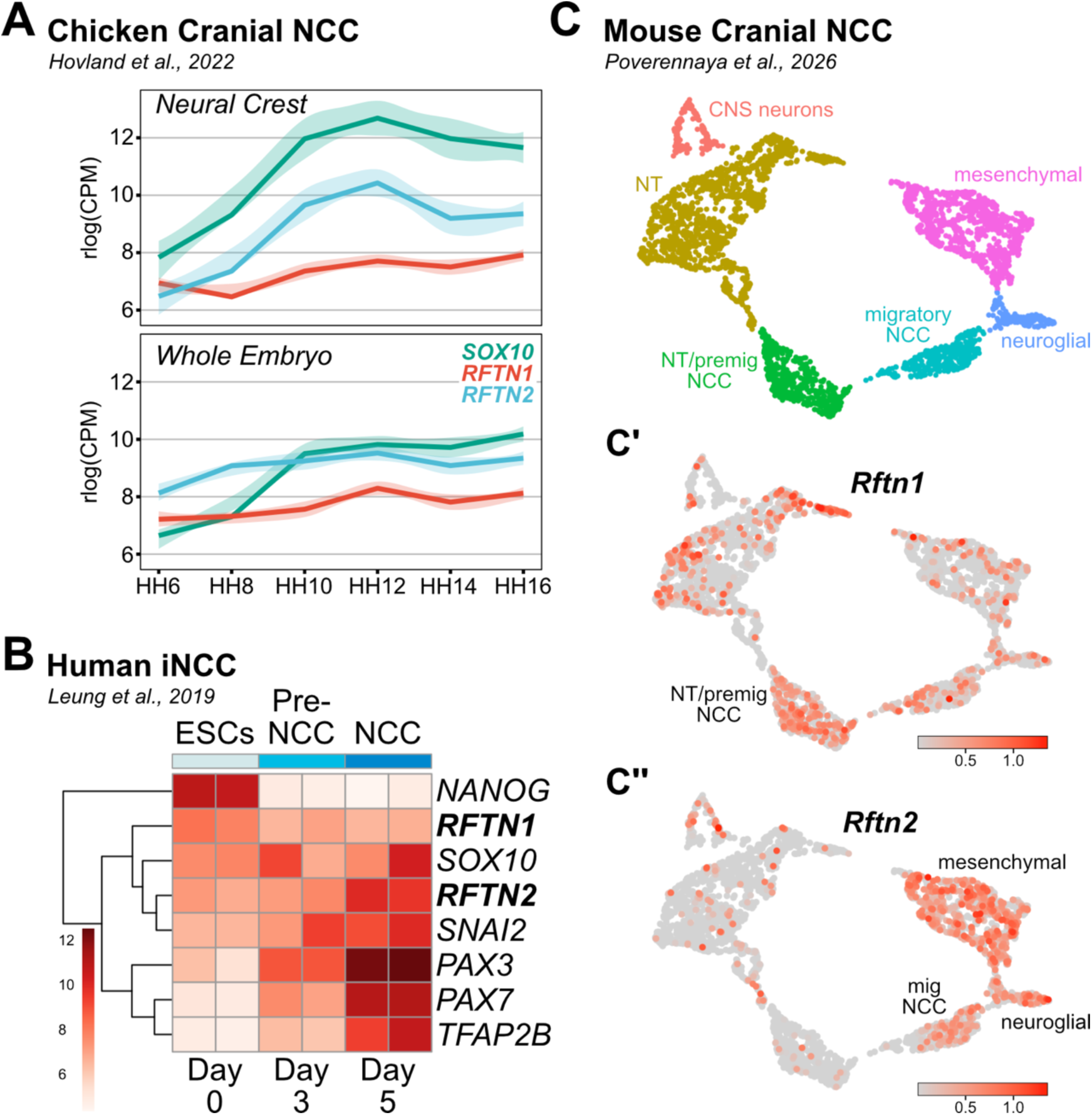
Raftlin genes show conserved expression patterns across amniotes. (A) Expression levels of *SOX10*, *RFTN1*, and *RFTN2* in regularized log-transformed counts per million (rlog(CPM)) results of bulk RNA sequencing from sorted chick neural crest cells (top panel) and the whole chick embryo (bottom panel); data from (Hovland et al., 2022). (B) Heatmap displaying variance-stabilizing transformation (VST)-normalized expression levels of indicated genes in human embryonic stem cells (ESCs), on Day 3 following Wnt activation to induce neural crest cell (NCC) fate (pre-NCC), and Day 5 of treatment representing NCC specification; data from (Leung et al., 2019). (C) UMAP cluster names (C) and indicated feature plots (C’, C’’) displaying single-cell transcriptomes from murine cranial neural crest sorted from *Wnt1-Cre/R26Tomato* transgenic mouse embryos at embryonic days E8.5, E9.5, and E10.5; data from (Poverennaya et al., 2026).

Next we examined bulk RNA sequencing from human neural crest cells differentiated from ESCs (Leung et al., 2019). Over a five-day induction period, ESCs show decreasing expression of pluripotency factors like *NANOG*, and increasing expression of markers of neural crest specification including *PAX3/7*, *TFAP2B*, *SNAI2*, and *SOX10* (Fig. 1B). Across this time course, *RFTN1* expression was detected at moderate levels across differentiation, with a mild reduction as ESCs start to differentiate. Conversely, *RFTN2* expression was detected within ESCs, and its expression strongly increased alongside bona fide markers of neural crest formation (Fig. 1B). Finally, we examined single-cell RNA sequencing results collected from sorted cranial neural crest cells from *Wnt1-Cre/R26Tomato* transgenic mouse embryos at embryonic days E8.5, E9.5, and E10.5 (Poverennaya et al., 2026). UMAP representation of these data capture neural tube (NT)-localized premigratory neural crest cells through their migration and fate restriction down either neuroglial or mesenchymal lineages (Fig. 1C). Feature plots displaying *Rftn1* expression show scattered expression throughout all cell clusters, with a slight enrichment in the NT/premigratory neural crest cluster (Fig. 1C’). Conversely, *Rftn2* shows sparse expression in the NT and premigratory neural crest, with clear expression biased to migratory and differentiating neural crest cell clusters (Fig. 1C’’). Together, these results suggest that expression of Raftlin-2, but not Raftlin, undergoes conserved upregulation during neural crest development across amniotes.

### Raftlin is broadly expressed throughout the embryo, with strong enrichment in the developing somites

To spatially resolve Raftlin gene family expression in wild type chick embryos, we designed hybridization chain reaction fluorescence *in situ* hybridization (HCR-FISH) probes specific to *RFTN1* and *RFTN2* and performed time course expression analyses alongside markers of early development primarily focused on neural crest and related tissues. We used third-generation HCR-FISH due to application of split probes for increased sequence specificity and reduce off-target signal (Choi et al., 2018), and initial experiments produced a distinct spatial expression pattern for each probe set (Fig. S1). To further test specificity, we also performed negative control experiments utilizing fluorescent hairpins without the corresponding Raftlin gene-specific probes. The results indicate that the observed labeling patterns are specific to Raftlin probe hybridization and not non-specific binding of labeling reagents (Fig. S1).

Upon whole mount analysis, the *RFTN1* expression pattern at HH10 appeared broadly ubiquitous with the most striking enrichment present in the neural tube at the level of the hindbrain, adjacent to the *PAX2-*positive otic placode, and in the most posterior paraxial mesoderm-derived somites; this somite-specific enrichment persisted through HH16 (Fig 2) and is consistent with *in situ* hybridization experiments labeling *Rftn* expression in the mouse tail bud (Chen et al., 2025). Next, we performed transverse sectioning to more precisely identify which tissues selectively upregulate *RFTN1* expression. In the head we observed broad, yet punctate staining across all major tissue populations including the neural tube (labeled by *PAX2* expression), cranial mesoderm and neural crest- (*TFAP2B*-expressing) and mesoderm-derived cranial mesenchyme, overlying non-neural ectoderm, and pharyngeal endoderm (Fig. 2A’). However, in more posterior sections at this stage, we observed similarly broad expression of *RFTN1* in all tissues, as well as enriched expression throughout the somite; this expression becomes more restricted to the ventromedial region of the somite in the more anterior, further developed somites (Fig. 2A’’-A’’’). This pattern is consistent with later stages of development at HH16, where we observed *RFTN1* expression broadly in the most posterior somites (Fig. 2B’’’’), restricted more ventromedially in the more anterior developing somites (Fig. 2B’’’), further restricted to the medial most sclerotome (Fig. 2B’’), and finally more diffuse, lower levels of expression in the most mature somites (Fig. 2B’). We also noted transient enrichment in *RFTN1* expression in the lateral plate mesoderm at HH10 (Fig.2 A’,A’’). Together these results are consistent with RNA-seq studies suggesting more ubiquitous and diffuse expression of *RFTN1* across development with limited neural crest cell enrichment (Fig. 1).

**Fig 2.**
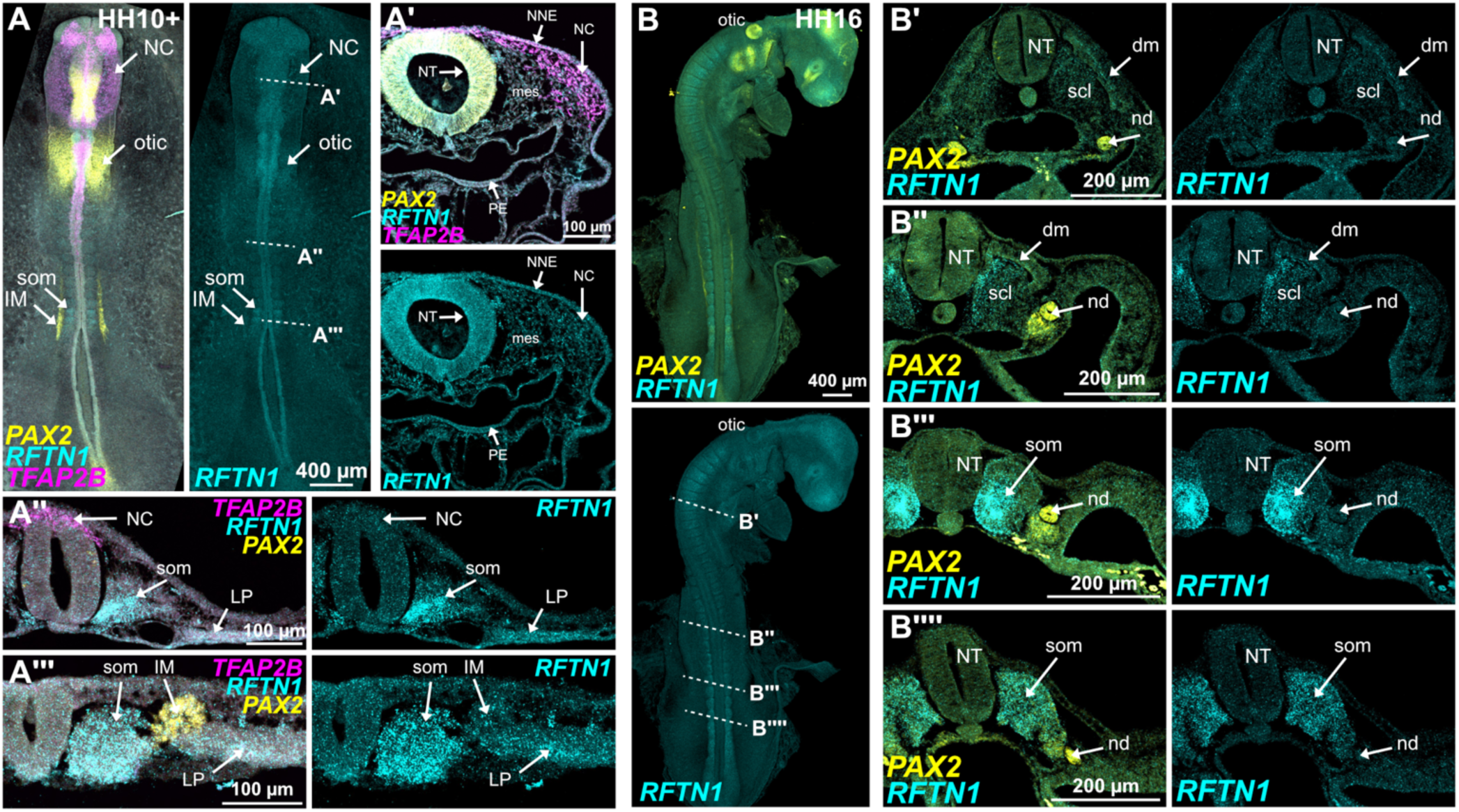
*RFTN1* is broadly expressed, with dynamic enrichment in developing somites. Wild type chick embryos were probed using HCR-FISH for expression of *RFTN1* alongside markers of neural crest cells (*TFAP2B*), and midbrain/hindbrain neural tube, otic placode, and intermediate mesoderm (*PAX2*) at the indicated stages. Transverse sections of wholemount embryos at axial positions indicated by dashed lines (A’-A’’’, B’-B’’’’). otic, otic placode; NC, neural crest; mes, cranial mesenchyme; NNE, non-neural ectoderm; PE, pharyngeal endoderm; som, somite; IM, intermediate mesoderm; LP, lateral plate mesoderm; nd, nephric duct; NT, neural tube; dm, dermamyotome; scl, sclerotome.

The expression pattern in the most posterior somites of HH10 and HH16 embryos raise the possibility that *RFTN1* plays an important role during early somitogenesis. Our results showed *RFTN1* expression to initially be highest in the newly epithelialized somites; however, after the ventromedial sclerotome undergoes epithelial-to-mesenchymal transition (EMT) and separates from the dermomyotome, this polarized enrichment is lost (Fig. 2B’’-B’’’). Interestingly, *RFTN1* expression in the ventromedial quadrant of the somite is reminiscent of the expression pattern of *PAX1* (Cauthen et al., 2001; Draga and Scaal, 2024). The transcription factor *PAX1* is essential for sclerotome development and its expression is induced by signaling molecules like Sonic hedgehog from neighboring tissues (Draga and Scaal, 2024; Fan and Tessier-Lavigne, 1994; Wallin et al., 1994). This raises the intriguing possibility that Raftlin may serve as a regulator of cell signaling events, such as through the Sonic hedgehog pathway, to facilitate sclerotome development. Future experiments will be required to test this hypothesis and assess if Raftlin works together with or upstream from *PAX1* expression during somite development.

### Raftlin-2 expression is enriched in neural crest cells during specification and early migration

We next sought to define the spatiotemporal pattern of *RFTN2* expression. Given evidence from RNA-sequencing studies that suggest enriched *RFTN2* expression in neural crest cell populations (Fig. 1)(Piacentino et al., 2022), we probed for *RFTN2* alongside *PAX7* to mark the induced neural crest cells within the neural plate border (Basch et al., 2006; Stundl et al., 2026). We did not detect appreciable *RFTN2* expression prior to neurulation within the well-defined *PAX7-*positive neural plate border at HH7 (Fig 3A). However as neurulation proceeds, specified neural crest cells at HH8 express *TFAP2B* (Rothstein and Simoes-Costa, 2020), at which point we observed the onset of *RFTN2* expression in the dorsal neural folds, and increasing expression the cranial mesoderm (Fig. 3B). *RFTN2* expression was expanded by HH9, detectable in nearly all *TFAP2B*-positive neural crest cells (Fig 3C-C’). Transverse sectioning confirms this pattern, revealing *RFTN2* signal to be most enriched in neural crest cells, and present at lower levels in the adjacent cranial mesoderm (Fig. 3C’’).

**Fig 3.**
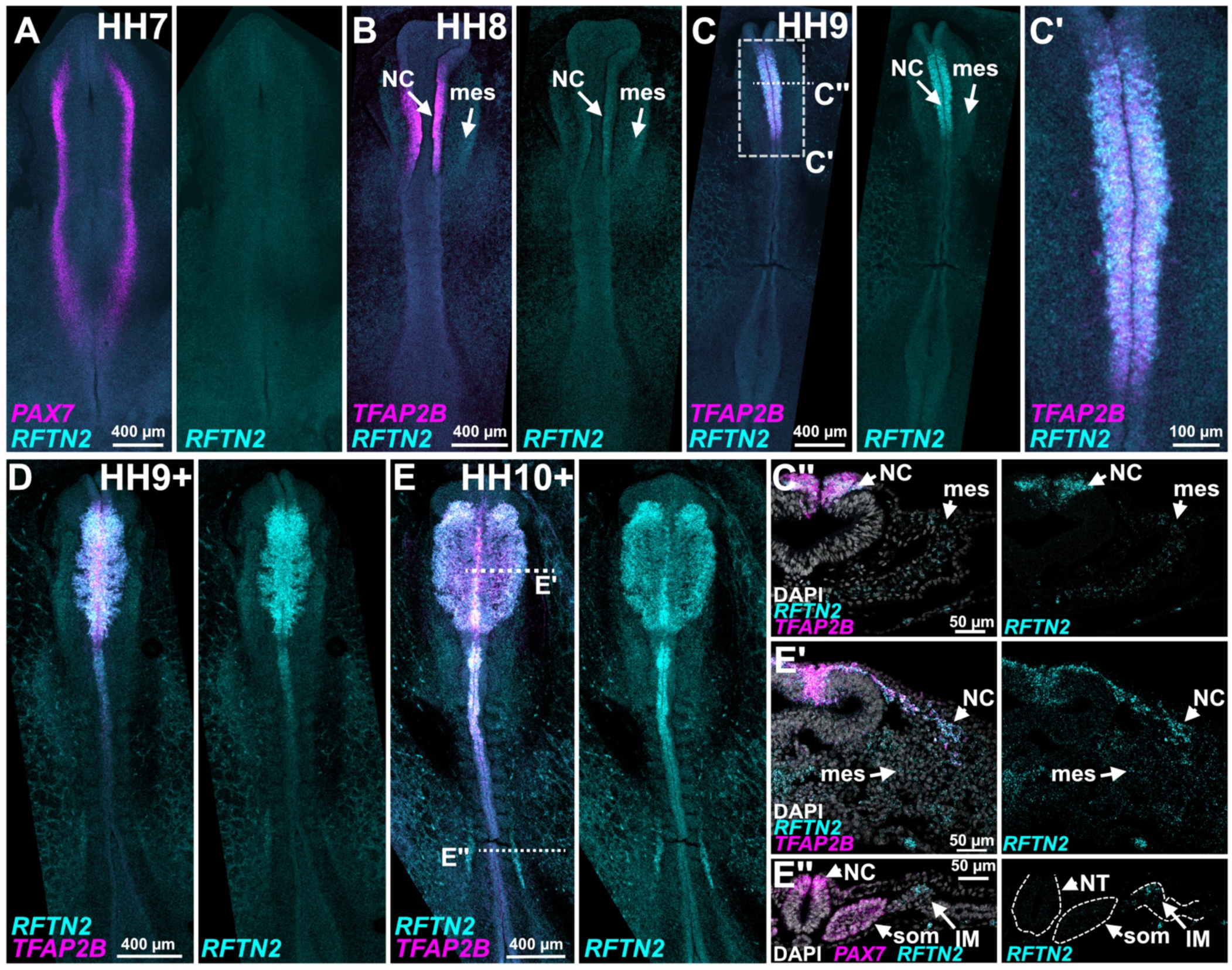
*RFTN2* is expressed in neural crest cells during specification and early migration. Wild type chick embryos were probed using HCR-FISH for expression of *RFTN2* and markers of neural crest cell development (*PAX7* and *TFAP2B*) at the indicated stages. Transverse sections of wholemount embryos at the axial levels indicated by dashed lines (C’, E’, E’’). Regions of higher magnification are designated by dashed boxes. NC, neural crest; mes, cranial mesenchyme; som, somite; IM, intermediate mesoderm, NT, neural tube.

Since *RFTN2* appeared to be upregulated in the neural crest after onset of markers of specification, it is unlikely that Raftlin-2 is required for neural crest cell induction or specification. However, given the upregulation of *RFTN2* expression after that of *TFAP2B*, it is possible that *RFTN2* is a target of the specification gene regulatory network and may contribute to maintaining specification and/or for neural crest cell survival before EMT and migration. Importantly, at this stage of development Wnt signaling remains an important facilitator of specification by activating expression of transcription factors like *FOXD3* through the effector protein Axud1 (Simões-Costa et al., 2015). In zebrafish embryos, Wnt signaling relies on the localization of the GPI anchored protein Lypd6 to lipid raft domains to facilitate the appropriate axis patterning (Özhan et al., 2013). Thus, it is possible that Raftlin-2 acts to advance neural crest development by facilitating lipid raft organization and promoting Wnt signaling.

We next probed *RFTN2* expression during early neural crest cell migration. *RFTN2* expression is maintained in migratory neural crest cells at stages HH9+ through HH10+ as these cells undergo and complete EMT (Fig. 3D-E). By transverse sectioning we confirmed robust *RFTN2* and *TFAP2B* expression in delaminating and migrating neural crest cells and found that *RFTN2* is expressed throughout the cranial mesoderm, albeit at a qualitatively lower level (Fig. 3E’). Transverse sections through the most posterior somites also reveal expression of *RFTN2* in the intermediate mesoderm (Fig. 3E’’), but not in the somites as was seen for *RFTN1* (Fig. 2). Since the intermediate mesoderm is the origin of nephric tissues and embryonic kidney (James and Schultheiss, 2003), this pattern suggests a possible role for Raftlin-2 in early kidney development.

As neural crest cells undergo EMT and delaminate, their migration is directed in response to chemotactic gradients, following chemoattractants such as vascular endothelial growth factor (VEGF) and stromal cell-derived factor 1 (SDF1) (Escot et al., 2013; McLennan et al., 2010; Olesnicky Killian et al., 2009; Shellard and Mayor, 2019; Wiszniak et al., 2015). Raftlin regulates VEGFR dependent migration in endothelial cells (Bayliss et al., 2020), raising the possibility that similar mechanisms mediated by Raftlin-2 may facilitate neural crest cell migration in response to VEGF. Additionally, the SDF-1 receptor, CXCR4, localizes to lipid rafts in hematopoietic stem cells where it enhances SDF-1 mediated migration (Wysoczynski et al., 2005). This supports the idea that lipid rafts, and potentially lipid raft organizing proteins, are important for cell signaling and migration in neural crest cells. Further, *RFTN2* expression present in the cranial mesoderm may reflect regulation of signaling activities between neural crest cells and the paraxial mesoderm in the head. For example, cranial neural crest cells secrete inhibitors of Wnt and BMP signaling that are important for cranial myogenesis (Tzahor et al., 2003) and mesoderm cells will briefly migrate with invading neural crest cells (McKinney et al., 2020). Shared signaling pathways and migration could indicate a common role for *RFTN2* in these two tissues.

### Raftlin-2 expression is downregulated during late neural crest cell migration into the pharyngeal arches

After observing robust expression at early stages of neural crest cell migration, we then probed *RFTN2* at later stages of migration and development. By HH13, *RFTN2* expression in the mesencephalic neural crest was reduced (Fig. 4A). However, the neural crest cells in the caudal hindbrain streams from rhombomeres r4 and r6 displayed strong *RFTN2* and *TFAP2B* expression as they migrated around the otic vesicle (Fig. 4A’). This is consistent with our previous observations where *RFTN2* is enriched during the earlier ventral-ward migration of the more rostral mesencephalic neural crest cells at HH10 (Fig. 3D-E). *RFTN2* expression also persisted in the most posterior intermediate mesoderm at HH13, while abundant *PAX2* expression was observed over a broader territory stretching anterior past the domain of *RFTN2* expression (Fig. 4A’’), suggesting that potential roles for Raftlin-2 in pronephros development may be transient. By HH17, *RFTN2* expression appears more diminished across the embryo in whole mount (Fig. 4B). Transverse sections through the head reveal clear expression of *TFAP2B* and *PAX2* in the pharyngeal arch ectoderm (Fig. 4B’), in tissues producing the cranial epidermis and epibranchial placodes, respectively (Baker et al., 2008; Van Otterloo et al., 2022; Washausen and Knabe, 2019). Notably, we observed slightly elevated *RFTN2* expression in the neural crest-derived mesenchyme of pharyngeal arch 1 and 2, which co-expressed low levels of *TFAP2B*, compared with the more medial cranial mesoderm-derived mesenchyme (Fig. 4B’). This neural crest population will contribute critical craniofacial skeletal elements including the mandible and hyoid (Frisdal and Trainor, 2014), thus, decreasing expression of *RFTN2* in the neural crest derived mesenchyme may suggest that its downregulation is required to permit ectomesenchymal differentiation.

**Fig 4.**
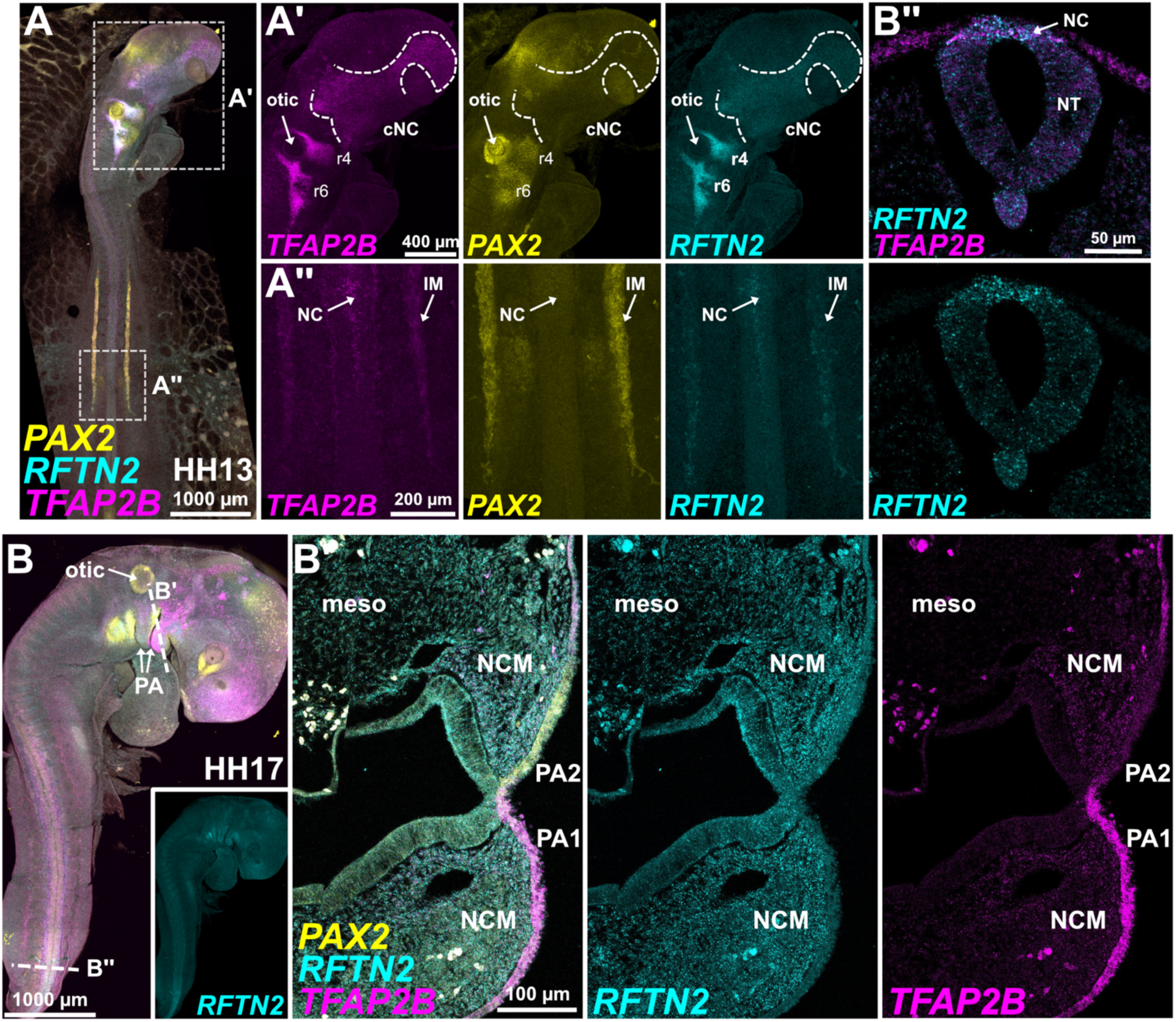
*RFTN2* expression is downregulated in late neural crest cell migration and maintained at a lower level in cranial neural crest-derived mesenchyme. Wild type embryos were probed using HCR-FISH for *RFTN2*, *TFAP2B*, and *PAX2* at the indicated stages. Transverse sections are indicated by dashed lines (B’-B’’). Regions of higher magnification are indicated by dashed boxes. (A’) Dashed line highlights the midbrain neural crest region. cNC, cranial neural crest; r4, rhombomere stream 4; r6, rhombomere stream 6; NC, neural crest; IM, intermediate mesoderm; NT, neural tube; PA, pharyngeal arch; meso, mesoderm; NCM, neural crest derived mesenchyme.

Consistent with *RFTN2* patterns of expression seen at earlier stages, trunk neural crest cells at HH13 and HH17 showed higher expression of *RFTN2* at the onset of migration (Fig. 4B’’). Since this expression appears downregulated as neural crest cells complete their migration into the pharyngeal arches of the face, this suggests the function of Raftlin-2 in the neural crest may be most closely linked to EMT and migration. Interestingly, given its expression pattern in the intermediate and cranial mesoderm, a role for *RFTN2* in cell migration could be utilized by multiple tissues. The intermediate mesoderm contributes to the nephric duct, and undergoes extension by collective migration starting at HH10 where leader cells migrate towards an FGF8 gradient produced in the tail bud, depositing cells to epithelialize along their migratory path (Atsuta and Takahashi, 2015; Soueid-Baumgarten et al., 2014). *RFTN2* expression is enriched in the most posterior portion of the intermediate mesoderm domain marked by *PAX2* at HH13 (Fig. 4A’’); this pattern reflects the location of the nephric duct leader cells as they actively migrate (Atsuta and Takahashi, 2015). Thus, *RFTN2* may be acting in the neural crest, cranial mesoderm, and intermediate mesoderm populations to modulate lipid raft activity to mediate migratory signaling events in each of these tissues.

## Conclusions

Our analysis has revealed that *RFTN1* and *RFTN2* have dynamic and specific expression patterns throughout early embryonic development. Using transcriptomic data sets, we demonstrated that these expression patterns are similar across vertebrate species: *RFTN1* is expressed lowly and broadly, while RFTN2 is upregulated during neural crest cell development. By probing *RFTN1* and *RFTN2* expression in the avian embryo, we showed that this pattern holds true *in vivo*. We observed sparse and broad *RFTN1* across the embryo with enrichment in somites, suggesting that *RFTN1* may play a role in somitic development. However, we found that *RFTN2* expression in neural crest tissues is dynamic with an increase in expression after the onset of specification that is maintained during early migration before being downregulated as these cells complete their migration. Our investigation also revealed *RFTN2* expression in the intermediate and cranial mesoderm which like neural crest cells have migratory abilities, suggesting that *RFTN2* may have specific roles in cell migration. These tissues must interpret signaling cues from their environments, implicating the Raftlin proteins as facilitators of cell signaling during the development of numerous embryonic tissues. These results call for more extensive *in vivo* functional studies to determine the specific and redundant roles of these proteins during development. Importantly, this area of inquiry presents an opportunity to inform our understanding of lipid raft functions during *in vivo* development.

## Acknowledgments

We would like to thank all the members of the Piacentino Lab for helpful discussions and specifically Alexander Pomper for contributions to protocol development. We would also like to thank Dr. Matteo Fabbri for help with anatomical identification. This work was supported by the National Institutes of Health of Dental and Craniofacial Research awards R00DE029240 (M.L.P.) and F31DE035770 (M.P.J.), and the National Institute of General Medical Sciences award R35GM90111815 (M.L.P.).

## Conflicts of interest

The authors have no conflicts of interest to declare.

## Author Contributions

Conceptualization: M.P.J. and M.L.P.

Experiment design: M.P.J., I.G., and M.L.P.

Experimentation: M.P.J., and I.G.

Data analysis: M.P.J., I.G., and M.L.P.

Data interpretation: M.P.J., I.G., and M.L.P.

Manuscript preparation: M.P.J. and M.L.P.

Manuscript editing: M.L.P.

## Materials and Methods

### Sequence analysis

Bulk and single cell RNA sequencing analysis was performed using previously published datasets. Briefly, regularized log-transformed counts per million (rlog(CPM)) were plotted from chick embryo sequencing data using a publicly available R Shiny app (http://ash274.shinyapps.io/RNA-Seq_App/)(Hovland et al., 2022). Human ESC-derived neural crest cell sequencing data was downloaded from the Gene Expression Omnibus (GSE125145, (Leung et al., 2019)). Reads were trimmed using fastp to remove low quality reads and overrepresented sequences (Chen et al., 2018). Reads were then mapped to a human reference genome (GRCh38 version 41) using STAR (Dobin et al., 2013) and count matrices were made using featureCounts (Liao et al., 2014). Variance-stabilized transformation (VST) was performed using DESeq2 (Love et al., 2014), and VST normalized expression values were plotted as heatmaps using Pheatmap in R (Kolde, 2015). For single cell RNA sequencing analysis, a processed An Anndata object describing gene expression from sorted *Wnt1-Cre/R26Tomato* transgenic mouse embryos was downloaded (Poverennaya et al., 2026), converted into a Seurat object using SeuratDisk, and custom feature plots were then generated using Seurat in R (Satija et al., 2015). Sequence analysis code was produced using Claude Sonnet v4.5 (Anthropic) and all code was manually reviewed, tested, revised, and verified by the authors. R code used for plotting human and mouse data are available (https://github.com/PiacentinoLab/2026_Jenne_RFTN_Expression).

### Chicken embryo collection

Fertilized white leghorn chicken eggs were purchased from commercial sources (University of Connecticut) and cultured at 38°C in humidified incubators until embryos reached the desired Hamburger Hamilton stage. At this point, embryos were dissected on filter paper into 1x Ringer’s solution, fixed for one hour at room temperature in 4% paraformaldehyde (PFA) in phosphate buffered saline (PBS,) rinsed in PBS with 0.1% Tween 20 (PBST), and dehydrated in a methanol/PBST series before storage in 100% methanol at −80°C.

### Hybridization chain reaction fluorescent in situ hybridization

Generation 3 hybridization chain reaction fluorescent *in situ* hybridization (HCR-FISH) labeling kits were purchased from Molecular Instruments (Choi et al., 2018). Probe sets were purchased directly from Molecular Instruments or custom designed using published pipelines and ordered as oPools (Integrated DNA Technologies)(Kuehn et al., 2022). Probe sets used in this study include *RFTN2* (B3 initiator), *RFTN1* (B5), *PAX7* (B2), *TFAP2B* (B4), *PAX2* (B2) and were detected using appropriate hairpin labeled with AlexaFluor488, AlexaFluor546, AlexaFluor647. Experiments were performed following manufacturer’s instructions. Briefly, dehydrated embryos were rehydrated in a methanol/PBST series, permeabilized in a Proteinase K (New England Biolabs, #P8107S) 0.4U/mL PBST solution for 1-2 minutes, optimized to stage/size of embryos, and post fixed for 20 minutes in 4% PFA in PBS. Embryos were washed twice in PBST, twice in 50% PBST/50% 5x saline sodium citrate 0.1% Tween 20 (SSCT), then twice for in 5x SSCT, each for five minutes at room temperature. Embryos were incubated in warmed probe hybridization buffer for 30 minutes at 37°C with rocking, then incubated overnight in probe hybridization buffer at 1:250 probe dilution at 37°C with rocking. Embryos were washed four times for 15 minutes in prewarmed (42 °C) probe wash buffer at 37°C, then washed twice for five minutes in 5x SSCT, then incubated in room temperature hybridization buffer for 30 minutes. Hairpins were snap-cooled separately by heating for 90 seconds at 95°C, then cooling at room temperature for 30 minutes. Embryos were incubated in a solution of amplification buffer at a 1:50 hairpin dilution overnight in the dark. Finally, embryos were washed 2 times for 5 minutes in 5x SSCT, post-fixed in 4% PFA in 5xSSC 0.1% Tween for 20 minutes at room temperature, washed 2 times for 5 minutes in 5x SSCT, washed twice for 30 minutes in 5x SSCT with 1:1000 dilution of 1 mg/mL DAPI on the first wash, 5 minutes in 5x SSCT.

### Sectioning

To generate transverse sections after whole mount HCR-FISH, labeled embryos were washed for five minutes in 50% PBST/50% 5x SSCT, washed five minutes in PBST, incubated in 5% sucrose in PBS for 15 minutes at room temperature, 15% sucrose in PBS overnight at 4°C, then 7.5% gelatin at 38°C overnight. Embryos were embedded in molds and flash frozen in liquid nitrogen before −80°C storage. Embryos were subsequently cryosectioned at a thickness of 18-20 µm.

### Imaging

Wholemount and transverse section imaging was performed using a Zeiss LSM900 Confocal with Airyscan2 Detector. Fluorescent wholemount and transverse section images were collected as Z stacks. Images were prepared for display using Fiji, and are presented as median-filtered maximum intensity projections.

## Supplementary Figures

**Fig S1.**
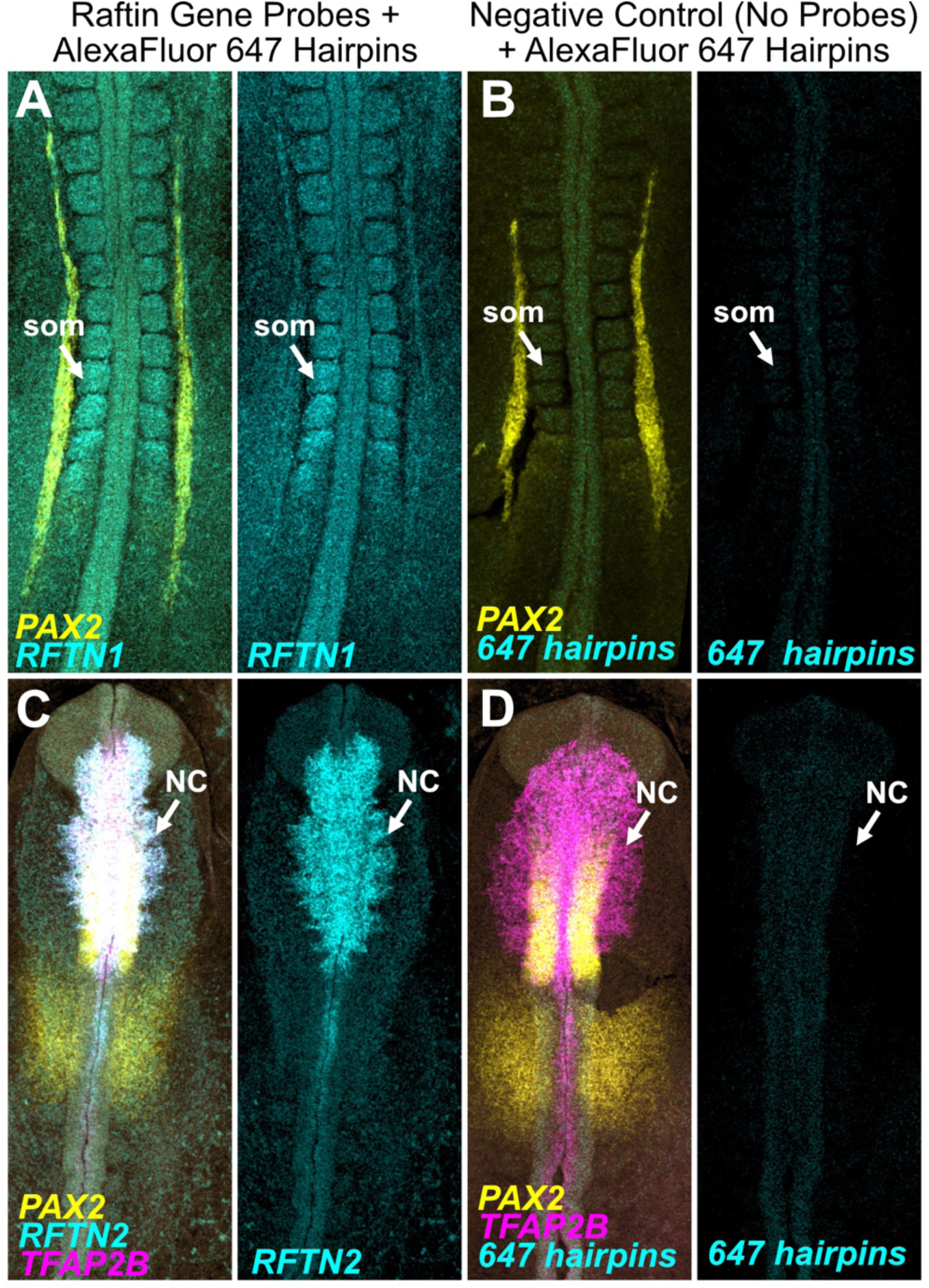
Labeling patterns for *RFTN1* and *RFTN2* HCR-FISH are not due to non-specific hairpin binding. Wild type embryos were processed for HCR-FISH to probe for expression of *RFTN1* and *PAX2* (A) or *RFTN2*, *TFAP2B*, and *PAX2* (C). Raftlin gene probes were detected using AlexaFluor 647-labeled hairpin amplification. Negative control experiments were performed using stage-matched wild type embryos, processed for HCR-FISH as above including AlexaFluor 647-labeled hairpins, but without the corresponding Raftlin gene probes (B, D). som, somite; NC, neural crest.

## Notes

https://github.com/PiacentinoLab/2026_Jenne_RFTN_Expression

### Competing Interest Statement

The authors have declared no competing interest.

